# Dynamic Compression of Spheroid-Laden Alginate Granular Composites Induces Hypertrophic Chondrocyte Phenotype

**DOI:** 10.64898/2026.03.14.711819

**Authors:** David H. Ramos-Rodriguez, Andrea C. Filler, Sheetal R. Palle, Shierly W. Fok, Erika E. Wheeler, J. Kent Leach

## Abstract

Hypertrophic cartilage is a promising bone repair strategy by producing a mineralizable matrix that transitions to bone through endochondral ossification. Current approaches require large cell numbers and costly recombinant factors to induce chondrogenesis. Here, we developed a composite granular scaffold using photocrosslinkable alginate microgels, cell-secreted decellularized extracellular matrix (dECM), and mesenchymal stromal cell (MSC) spheroids under dynamic compressive loading for hypertrophic cartilage formation. Incorporation of dECM into MSC spheroids enhanced expression of chondrogenic markers and supported the hypertrophic phenotype, evidenced by increased VEGFA and SPP1 expression and ALP activity. Dynamic loading further increased spheroid sprouting and scaffold mineralization. Histology confirmed mature hypertrophic cartilage conducive to bone formation. Upregulation of hypertrophic and osteogenic markers was associated with YAP1 activation, linking compressive loading to mechanotransduction to drive hypertrophic cartilage formation. These results demonstrate that dynamic compressive loading, cell aggregates, and scaffold granular macroporosity synergistically yield hypertrophic cartilage.

## Introduction

Bone defects resulting from trauma, tumor resection, or congenital abnormalities represent a significant clinical challenge.(1) Autologous bone graft remains the clinical gold standard, but current grafting approaches are often hindered by donor site morbidity, limited tissue availability, and poor integration, particularly in large defects. Although tissue engineering strategies offer a promising alternative to bone grafting, conventional approaches that rely on direct bone formation frequently fail to achieve sufficient graft stability, vascularization, and long-term functionality. Leveraging the body’s natural endochondral ossification (EO) process, in which a transient cartilage template orchestrates vascular invasion and bone regeneration, offers a biologically inspired route to improve the translation of regenerative therapies for mineralized tissue.(2)

Endochondral bone formation from mesenchymal stromal cells (MSCs) and biomaterial scaffolds has been reported,(3, 4) but consistent development of hypertrophic phenotype, structural stability, and controlled remodeling limit clinical translation. These hurdles are exacerbated by the necessity of large numbers of chondrocytes capable of undergoing hypertrophic differentiation and producing the essential ECM for construct volume. Therefore, developing a construct that reproduces the cellular and extracellular hallmarks of hypertrophic cartilage is critical to unlocking the translational potential of EO-based bone regeneration.

Granular hydrogels have emerged as versatile building blocks for engineering replacement tissues or promoting tissue formation due to increased modularity and precise control over cell–matrix interactions.(5–9) Their high surface-to-volume ratio facilitates efficient nutrient transport, while tunable biochemical and mechanical properties enable fine regulation of cell fate. Mechanical stimulation enhances chondrogenic and hypertrophic differentiation in bulk hydrogel systems by activating mechanotransduction pathways and cytoskeletal reorganization, ultimately driving matrix remodeling, collagen deposition, and mineralization.(10–12) However, the impact of mechanical loading on granular or microgel-based scaffolds remains largely unexplored. The porous and modular architecture of microgel scaffolds may allow microscale stress redistribution and localized mechanotransduction not achievable in bulk gels. While microgels have been fabricated from synthetic and natural polymers such as polyethylene glycol, hyaluronic acid, and gelatin(5), we fabricated granular hydrogels from alginate due to its tunable stiffness, degradability, ease of functionalization, and viscoelastic characteristics similar to native chondrogenic tissues.(13) These properties make alginate microgels particularly suitable for recreating dynamic cartilage microenvironments that guide chondrogenic and hypertrophic differentiation.

Chondrogenic and hypertrophic differentiation in vitro is induced by high concentrations of recombinant inductive factors to regulate cell fate. The extracellular matrix (ECM) plays an essential role in growth factor signaling by providing structural support and biochemical and biophysical signals that synergistically coordinate cell adhesion, proliferation, and lineage progression.(14, 15) As an alternative to decellularized tissues as a source of ECM, mesenchymal stromal cell (MSC)-secreted decellularized ECM (dECM) can be engineered as an on-demand, instructive template to guide cell fate with the potential for reduced immunogenicity.(16, 17) The interaction of cells with dECM promotes robust, tissue-specific differentiation without supraphysiologic growth factors by recapitulating native matrix composition, matrix-bound signals, and spatial motifs to guide differentiation and integration.(18–20)

To advance the field of hypertrophic cartilage formation, we hypothesize that composite scaffolds composed of alginate microgels and MSC spheroids can effectively promote hypertrophic differentiation when applying physiologically relevant mechanical compression and facilitate mineralized tissue formation through EO. Our goal is to provide chondrogenic and hypertrophic cues while simultaneously supplying a biomimetic matrix that supports mineralization for EO-driven bone regeneration.

## Results

### Decellularized ECM enhances chondrogenic priming and development of hypertrophic phenotype in MSC spheroids

Human bone marrow-derived MSC spheroids were cultured for 35 d to induce hypertrophic differentiation (Figure 1A). Building upon our previous findings that dECM enhances chondrogenic differentiation(19), we now show that its inclusion enhances responsiveness to hypertrophic cues. Our results suggest that dECM incorporation into MSC spheroids and culture for 14 d in chondrogenic media (CM) increased spheroid diameter by 1.9-fold (282 ± 37 µm vs 149 ± 21 µm, p<0.0001) (Figure 1B), increased DNA content per spheroid (898 ± 73 pg vs 365 ± 20 pg, p<0.0001) (Figure 1C), and upregulated secretion of sulfated glycosaminoglycans (sGAGs) by 2.5-fold (17 ± 3 pg vs 7 ± 3 pg, p<0.0001) (Figure 1D) compared to control spheroids. Increased production of sGAGs was accompanied by expression of chondrogenic genes for both spheroid groups (Figure 1E-F). We did not observe significant differences in SOX9 (p=0.13) and ACAN (p=0.5) expression despite increases in sGAG content. These data suggest that matrix synthesis was initiated earlier and matrix retention is enhanced or degradation is reduced within dECM-spheroids, which collectively support sustained preservation of a GAG-rich cartilage matrix. In agreement with sGAG quantification, Safranin-O staining and immunohistochemistry (IHC) revealed a higher expression of sGAGs and collagen II for dECM spheroids, respectively (Figure 1K). Collectively, these data highlight the potential of dECM to accelerate chondrogenic priming in MSC spheroids by increasing production and retention of cartilage-like matrix components.

**Figure 1.**
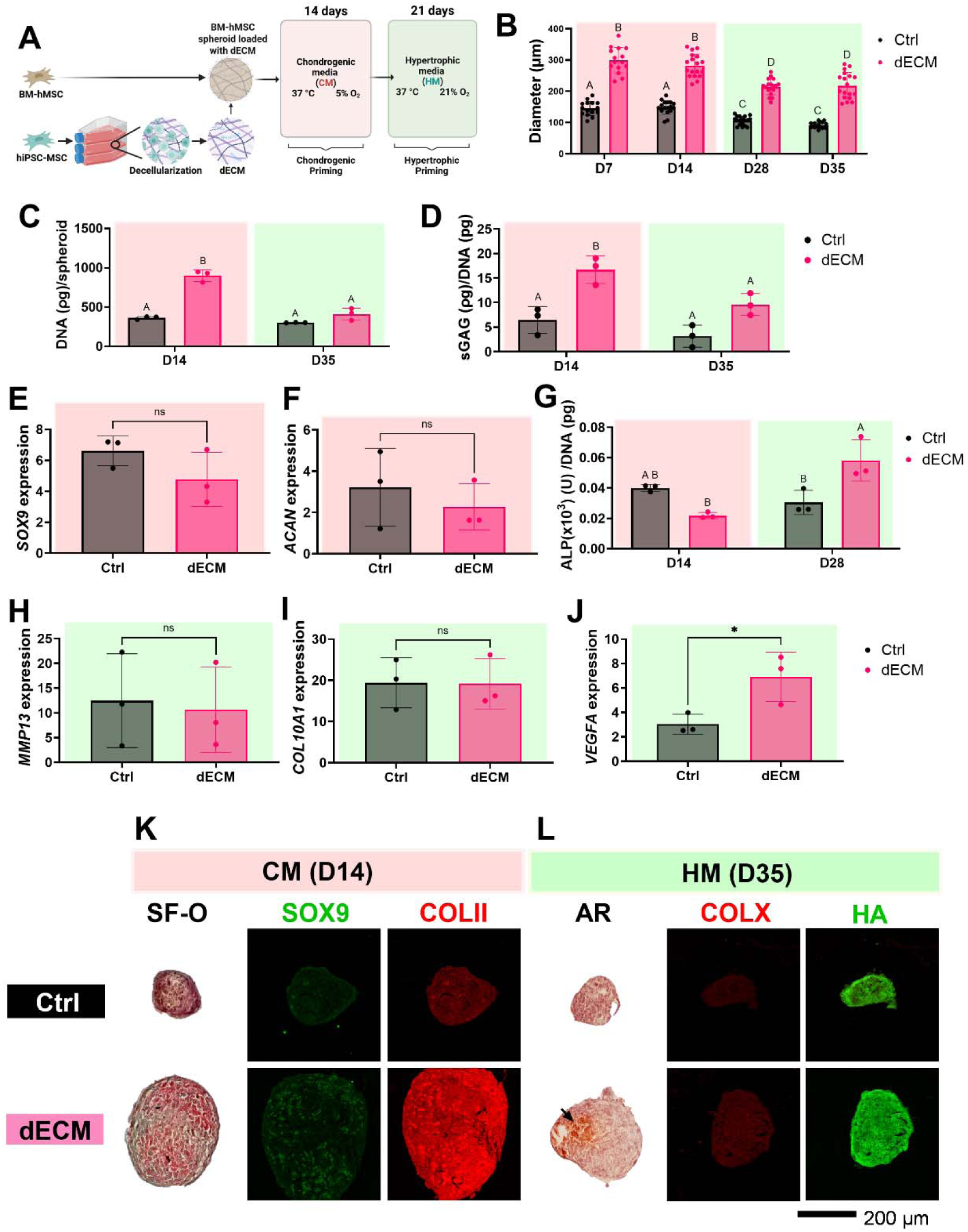
Introduction of dECM increases expression of chondrogenic markers by d14 and supports progression of hypertrophic phenotype in BM-MSC spheroids. **(A)** Schematic diagram of the fabrication of hypertrophic spheroids loaded with dECM. **(B)** Spheroid diameter (n>15); **(C)** DNA content; and **(D)** normalized sGAG content at different timepoints during chondrogenic (pink) and hypertrophic (green) conditioning. Expression of chondrogenic genes **(E)** *SOX9* and **(F)** *ACAN* after 14 d in CM. **(G)** ALP activity measured at d 14 and d 28 after chondrogenic and hypertrophic priming. Analysis of gene expression at d35 for hypertrophic genes **(H)** *MMP13***, (I)** *COL10A1*, and **(J)** *VEGFA*. Histological analysis for **(K)** chondrogenic and **(L)** hypertrophic markers. Scale bar is 200 µm. Bars with different letters are statistically different from each other. Data are mean ± standard deviation (n=3). *p<0.05. **Ctrl**, spheroids lacking dECM; **dECM**, spheroids loaded with dECM; **SF-O**, safranin-O; **COLII**, collagen type II; **AR**, alizarin red; **COLX**, collagen type 10; **HA**, hydroxyapatite.

After chondrogenic priming, dECM spheroids cultured for 21 d in hypertrophic media (HM) remained larger than control spheroids by 2.4-fold. Yet, when compared to spheroids in CM, we observed a consistent reduction in diameter (282 ± 36 µm vs 218 ± 42 µm; p<0.0001) and DNA content (898 ± 73.3 pg vs 412 ± 76.4 pg; p=0.14) for dECM-loaded spheroids by d 35 (Figure 1B-C). At the end of hypertrophic culture, we did not detect differences in sGAG content between control (3 ± 2 pg; p=0.19) or dECM (10 ± 2 pg; p=0.055) spheroids. However, we did observe a significant reduction in sGAG content for dECM spheroids after the hypertrophic culture period compared to sGAGs quantified after chondrogenic priming (p=0.035) (Figure 1D).

To evaluate the potential for tissue mineralization, we measured alkaline phosphatase (ALP) activity after 14 d in CM and additional 14 d in HM (Figure 1G). Compared to control spheroids, ALP activity of dECM spheroids was higher after 28 d (0.058 ± 0.01 U pg-1 vs 0.031 ± 0.01 U pg-1; p=0.01). The increase in ALP activity suggests a higher potential for tissue mineralization in dECM spheroids. Commitment to hypertrophic phenotype and early tissue mineralization was corroborated by increased collagen X staining, as well as high intensity Alizarin red and hydroxyapatite staining compared to control spheroids (Figure 1L).

Despite these trends, we did not detect differences in the expression of hypertrophic genes COL10A1 (p=0.96) and MMP13 (p=0.86) after 35 d in culture (Figure 1H-I). Nonetheless, upregulation of VEGFA in dECM spheroids (p=0.03) suggests higher commitment to hypertrophic phenotype compared to control spheroids (Figure 1J). Altogether, these results demonstrate the capacity of dECM to improve response of chondrogenic spheroids to hypertrophic cues. The increase in ALP activity, coupled with the upregulation of hypertrophic marker VEGFA, suggests greater potential for tissue mineralization in dECM spheroids. The presence of hydroxyapatite in dECM spheroids further supports their potential to serve as building blocks that support EO.

### Alginate microgels exhibit morphological homogeneity

Alginate is a bioinert, cytocompatible polysaccharide with predictable control over stiffness and degradation when crosslinked into a non-soluble matrix. However, its lack of intrinsic adhesion sites requires integration of defined ligands (e.g., RGD) that facilitate cell attachment. Although the effect of RGD ligands in alginate gels is well established for chondrogenic(21) and osteogenic(22) differentiation, there are no studies that explore the effects of RGD density in the development of hypertrophic chondrocyte phenotype using granular scaffolds. To address this knowledge gap, alginate was fabricated into monodisperse microgels that were combined into granular scaffolds with interconnected pore space and high effective diffusivity. We fabricated microgels with low and high RGD content to decouple biophysical from bioactive effects. Alginate microgels were engineered with independently tunable stiffness by crosslinking with Ca-EDTA and annealing via methacrylation (Figure 2A). Microgel diameter for low (116 ± 5 µm) and high (115 ± 4 µm) RGD content alginate microgels was comparable (p=0.22) with a low coefficient of variation (CV) of 3.9% for low and 3.6% for high RGD, indicating a monodisperse population (Figure 2B-C). Both microgel groups exhibited normal diameter distributions (Figure 2D) and similar compressive moduli (1.5 ± 0.4 kPa for low RGD vs 1.9 ± 0.8 kPa for high RGD; p=0.5) (Figure 2E), confirming homogeneous and mechanically consistent microgel populations regardless of RGD content.

**Figure 2.**
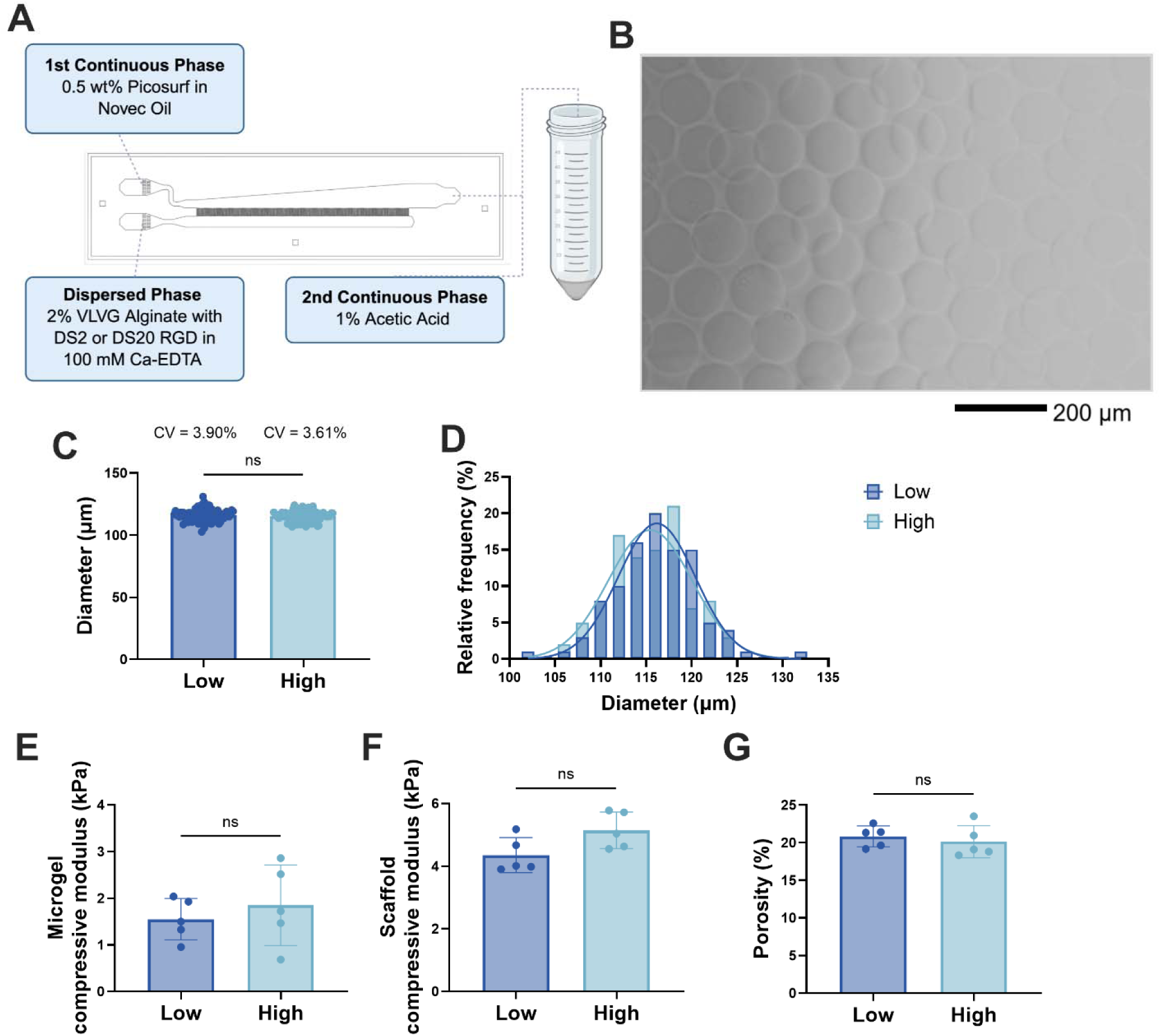
Characterization of alginate microgels and granular scaffolds. (**A**) Schematic of the fabrication of alginate microgels using a microfluidic approach. (**B**) Brightfield images of microgels post-crosslinking were used to quantify (**C**) microgel diameter and (**D**) size distribution (n=100). Scale bar is 200 µm. Similarities in (**E**) microgel compressive modulus, (**F**) scaffold compressive modulus, and (**G**) porosity revealed that RGD concentration did not affect biophysical properties. Data are mean ± standard deviation (n=5). **DS**, RGD degree of substitution; **CV**, coefficient of variation.

Successful photoannealing was confirmed when quantifying increased compressive modulus upon fabrication into an annealed scaffold, in agreement with observations from PEG-based annealed microgels.(23) We observed similar compressive moduli (4.4 ± 0.5 kPa for low RGD vs 5.1 ± 0.5 kPa for high RGD; p=0.059) (Figure 2F) and scaffold microporosity (20.8 ± 1.3% for low RGD vs 20.1 ± 2.1% for high RGD; p=0.5) (Figure 2G), regardless of RGD density (Figure S1). These findings indicate successful microgel annealing and uniform crosslinking when functionalizing alginate with different concentrations of RGD peptide. Overall, these data demonstrate that alginate microgels and resulting photo-annealed scaffolds exhibit consistent mechanical and structural properties regardless of RGD functionalization, representing an innovative substrate to support hypertrophic differentiation and EO.

### Hypertrophic differentiation is enhanced in composites containing dECM-loaded spheroids

Unlike bulk hydrogels possessing a nanoporous network, granular scaffolds provide an interconnected, cell-scale pore network with independently tunable properties.(5) This architecture enables efficient nutrient and waste transport, cell proliferation, cell migration, and differentiation.(24) To determine whether alginate granular scaffolds further promote hypertrophic chondrocyte differentiation and mineralization of MSC spheroids after chondrogenic priming, we assessed the expression of hypertrophic markers, focusing on the synergistic contribution of RGD ligand density and dECM incorporation (Figure 3A). After 21 d of hypertrophic priming, scaffolds exhibited comparable DNA content regardless of RGD concentration or inclusion of dECM (Figure 3B), indicating that these two variables had no differential effect on cell proliferation within the granular network.

**Figure 3.**
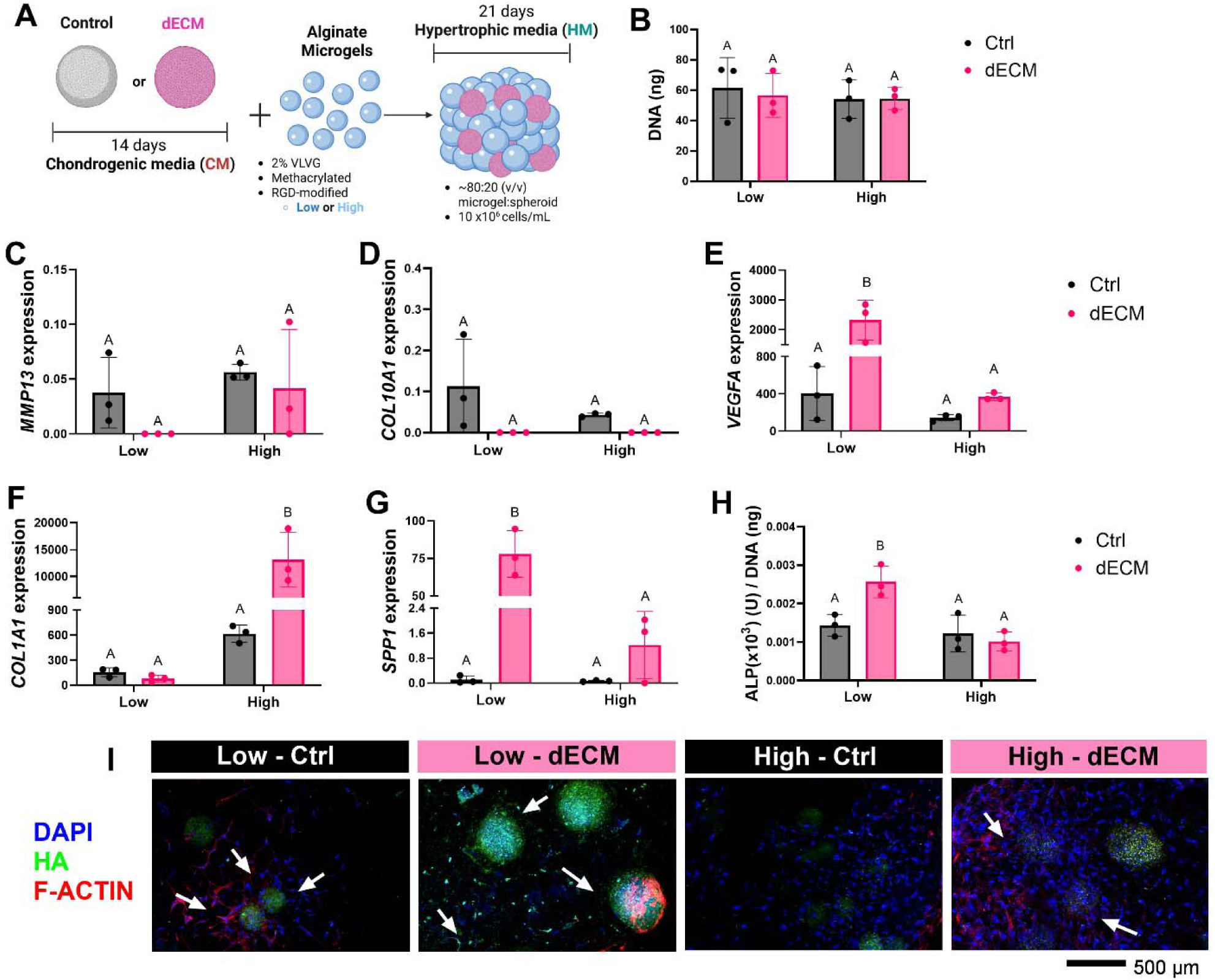
dECM-loaded spheroid-laden alginate granular scaffolds exhibit increased hypertrophic differentiation. (**A**) Schematic of fabrication of alginate granular scaffolds made from spheroids cultured in chondrogenic conditions for 14 d. Scaffolds were kept in static conditions and cultured in HM for 21 d. (**B**) DNA of scaffolds made from microgels with low or high RGD contents. Quantification of gene expression of hypertrophic markers (**C**) *MMP13,* (**D**) *COL10A1,* and (**E**) *VEGFA* across two levels of RGD covalent modification (low and high). Expression of osteogenic genes (**F**) *COL1A1* and (**G**) *SPP1* was used to evaluate early commitment to tissue mineralization. Osteoblastic differentiation was detected by (**H**) ALP activity and (**I**) HA staining. Scale bar is 500 µm. Bars with different letters are statistically different from each other. Data are mean ± standard deviation (n=3). **Ctrl**, spheroids without dECM; **dECM**, spheroids loaded with dECM; **SF**, safranin-O; **COLII**, collagen type II; **AR**, alizarin red; COLX, collagen type X; **HA**, hydroxyapatite.

Analysis of hypertrophic gene expression revealed a low, nonsignificant expression of MMP13 (Figure 3C) and COL10A1 (Figure 3D) for all groups, regardless of RGD content or inclusion of dECM-loaded spheroids. Despite low expression of hallmark hypertrophic genes, all scaffolds exhibited high VEGFA upregulation by d35 (Figure 3E). Notably, dECM-loaded spheroids in low RGD exhibited higher expression compared to control spheroids and spheroids in high RGD alginate (p<0.001 for all comparisons). These findings suggest that dECM elicits a proangiogenic response compatible with maturation of hypertrophic phenotype and signal endochondral progression.

We next assessed markers of osteogenesis in scaffolds undergoing hypertrophic culture in alginate microgels presenting low and high RGD content. The expression of osteogenic genes, including COL1A1 (Figure 3F) and SPP1 (Figure 3G), was largely increased when spheroids were loaded with dECM compared to control spheroids, regardless of RGD content, with exception of dECM-loaded spheroids in low RGD alginate scaffolds. Consistent with a shift towards mineralization, ALP activity (Figure 3H) was 1.85-fold higher at d 35 in low RGD dECM-loaded spheroids versus control spheroids of the same RGD content. Overall, dECM-loaded low RGD outperformed its high concentration counterpart (p=0.003) and both low (p<0.02) and high (p=0.009) RGD control scaffolds in ALP activity and SPP1 expression (p<0.001), reinforcing the combinatorial benefit of dECM presentation within granular architecture. Immunohistochemistry and detection of cell mineralization (Figure 3I) corroborated these findings at the microstructural level. F-actin staining revealed organized cytoskeletal fibers and spheroid protrusions in low and high RGD controls. Interestingly, the greater mineralization observed through hydroxyapatite staining in low RGD dECM-loaded scaffolds was accompanied by low spheroid sprouting, suggesting that continued aggregation may enhance mineralization, similar to previous work.(25)

Cell staining revealed distinct differences in osteogenic and hypertrophic marker expression among groups (Figure 4). Granular scaffolds containing dECM-loaded spheroids exhibited strong staining for osteocalcin (OCN) and collagen I (COL1), indicating enhanced osteogenic activity compared to control scaffolds. Notably, dECM-loaded spheroids cultured in high RGD scaffolds displayed pronounced cellular sprouting accompanied by reduced spheroid diameter, suggesting active-matrix remodeling and outward migration of hypertrophic chondrocytes. In contrast to control scaffolds, dECM-loaded scaffolds exhibited minimal collagen X staining, while showing robust expression of osteogenic markers. This pattern indicates progression beyond the hypertrophic phase toward a more advanced stage of endochondral maturation and matrix mineralization.

**Figure 4.**
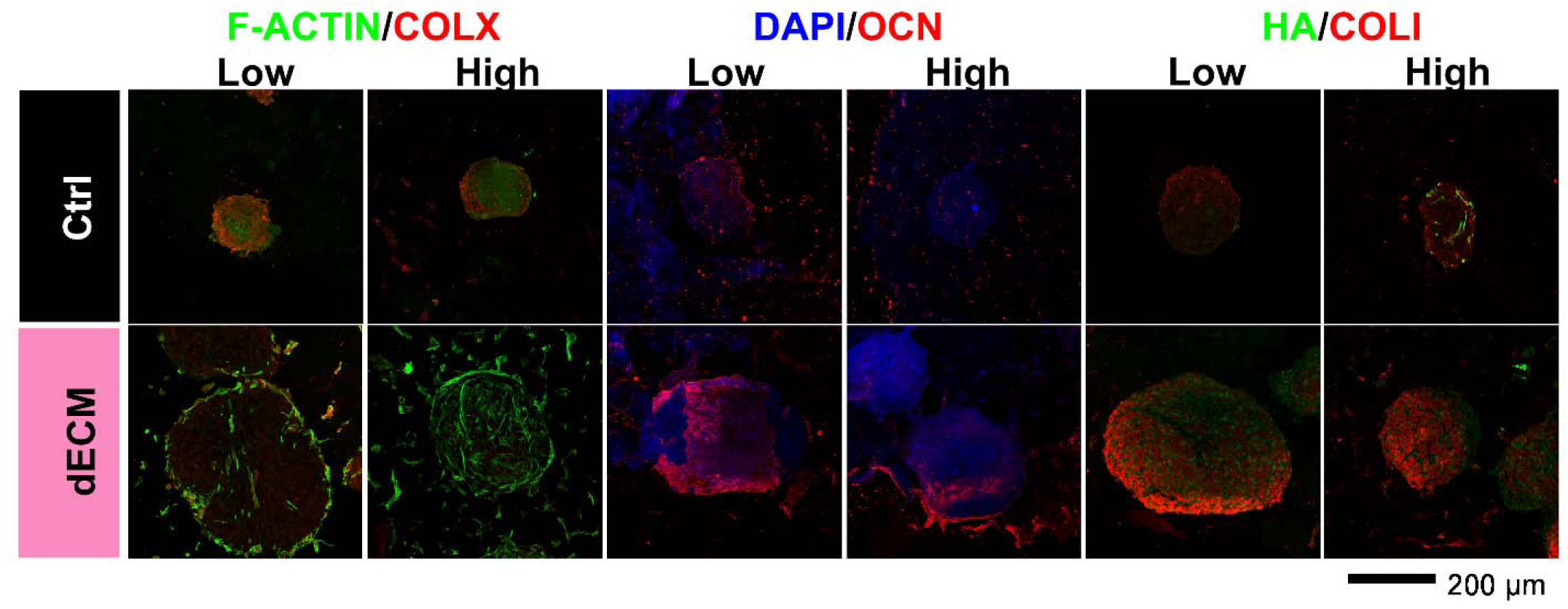
RGD content in annealed alginate microgel scaffolds and dECM inclusion improves progression toward the hypertrophic phenotype in MSC spheroids. Histological evaluation of dECM-loaded granular scaffolds at d 35. Representative images reveal differences in expression for F-actin, collagen X (COLX), cell nuclei (DAPI), osteocalcin (OCN), hydroxyapatite (HA), and collagen I (COLI) between control (Ctrl), dECM, low, and high RGD content groups. Scale bar is 200 µm.

Collectively, these data demonstrate that dECM-loaded spheroid-laden alginate granular scaffolds can drive hypertrophic phenotype while supporting angiogenic signaling and accelerating the osteogenic transition. We observed that RGD content modulated dECM spheroid response, as mineralization increased in the low-RGD scaffolds, indicating a synergistic effect between ligand density and dECM cues. However, low RGD scaffolds supported limited cell outgrowth within the construct.

### Dynamic compressive loading accelerates development of hypertrophic phenotype and enhanced mineralization

Mechanical stimulation within tissue engineered scaffolds is a key regulator of stem cell differentiation and can accelerate the development of functional tissue properties. Unlike bulk hydrogels, cells interact with granular hydrogels through localized microenvironments that enable dynamic interactions largely independent of the scaffold’s bulk mechanical behavior. These alginate microgels form a viscoelastic matrix that undergoes stress relaxation, a feature known to promote mechanotransducive signaling and lineage specification toward osteogenic or chondrogenic progenitors(26, 27). In this work, we investigated the synergistic effects of mechanical loading, RGD ligand density, and the incorporation of dECM into MSC spheroids on the development of a hypertrophic chondrocyte phenotype.

Our results demonstrate that a 5-day regimen of compressive loading applied to granular scaffolds, followed by 16 d of static culture (Figure 5A), did not diminish the structural integrity of the annealed constructs (Figure S2). Furthermore, compressive loading promoted spheroid sprouting and enhanced cytoskeletal organization, evidenced by increased F-actin filament expression (Figure 5B). This transient loading period appears sufficient to activate sustained mechanotransductive signaling, especially in dECM-loaded spheroids, likely mediated by integrin clustering and downstream actin polymerization. The resulting enhancement in cytoskeletal tension may facilitate YAP/TAZ nuclear translocation and transcriptional activation of genes associated with matrix remodeling and chondrogenic hypertrophy. In contrast, cells in high RGD control scaffolds were less responsive to mechanical stimulation, evidenced by unchanged cell outgrowth and cytoskeletal alignment, suggesting that excessive ligand density may restrict dynamic adhesion turnover required for optimal mechanosensing. We observed a marked decline in cellular infiltration and spheroid sprouting in low RGD control scaffolds by d 35 relative to d 21, indicative of cytoskeletal relaxation and diminished cell–matrix communication in the absence of mechanical cues. Collectively, these findings highlight the synergistic role of mechanical loading and dECM in regulating mechanotransduction pathways.

**Figure 5.**
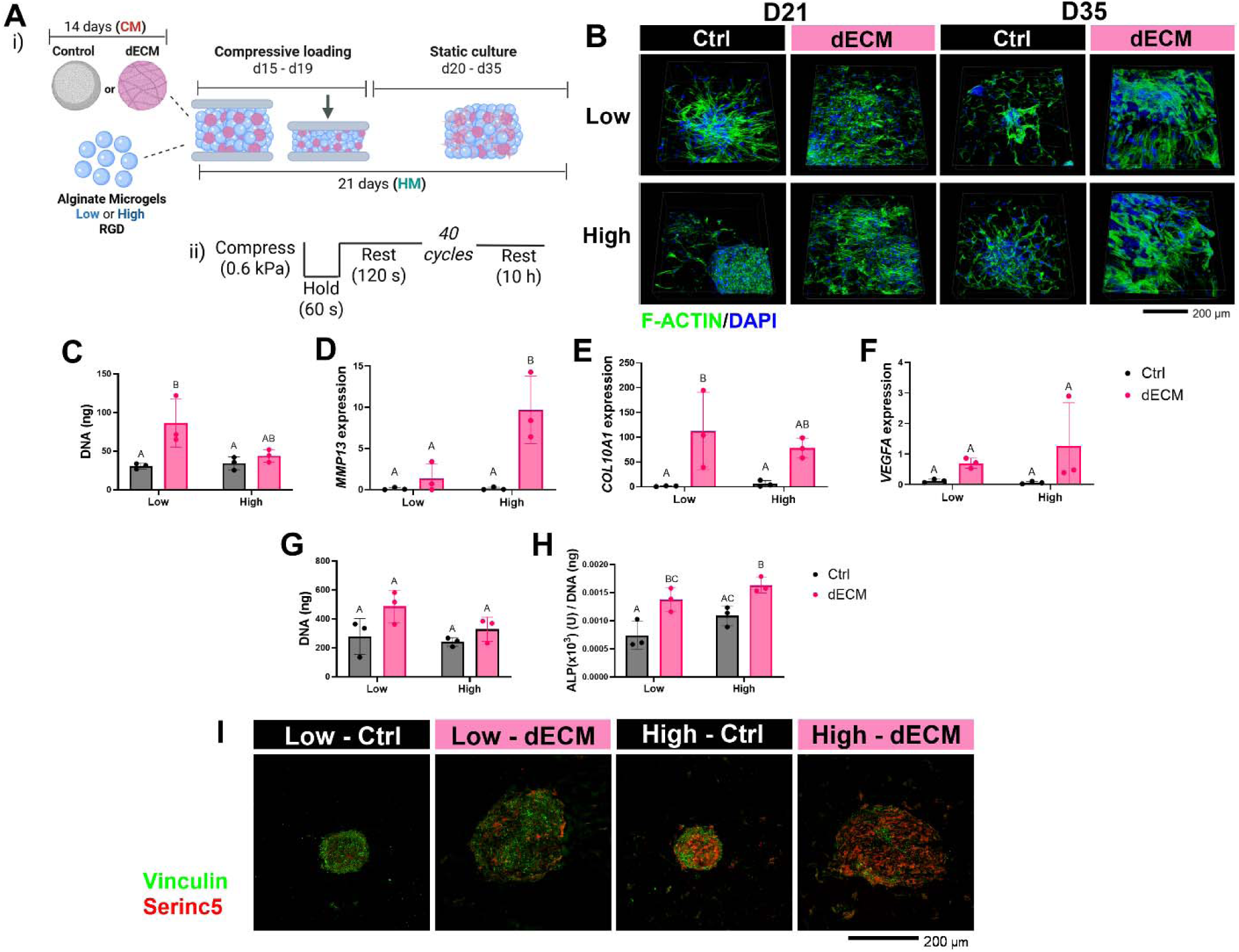
Dynamic mechanical compression promotes spheroid sprouting, cell proliferation, and upregulates hypertrophic markers on dECM-loaded granular composite scaffolds. (**A**) Schematic of fabrication of alginate granular spheroid-laden scaffolds cultured in chondrogenic conditions for 14 d, followed by dynamic compression for 5 d, and cultured in HM under static conditions until d 35. (**B**) Fluorescent images of spheroid-laden granular composite scaffolds at 21 and 35 d reveal differences in cytoskeleton architecture (F-actin; green) and cell proliferation (DAPI; blue). Scale bar is 200 µm. (**C**) Quantification of DNA content and gene expression after dynamic compression (d 21) of hypertrophic markers (**D**) *MMP13,* (**E**) *COL10A1,* (**F**) *VEGFA*. (**G**) DNA content and (**H**) ALP activity of granular scaffolds at d 35 made from low and high RGD-modified alginate microgels were quantified to characterize cell proliferation and early osteoblastic activity, respectively. (**I**) Serinc5 and vinculin staining were used to characterize progression of hypertrophic phenotype at d 35. Scale bar is 200 µm. Bars with different letters are statistically different from each other. Data are mean ± standard deviation (n=3). **Ctrl**, spheroids without dECM; **dECM**, spheroids loaded with dECM.

At d 21 (24 h after compressive loading), DNA content was ∼2-fold higher for low RGD dECM scaffolds (86 ± 31 ng) compared to low RGD control and high RGD scaffolds, indicating greater cell proliferation (Figure 5C). Analysis of gene expression at d 21 revealed upregulation of MMP13 and COL10A1 in mechanically stimulated dECM-loaded MSC spheroid composites, indicating enhanced matrix remodeling and progression toward hypertrophic maturation compared to the control group (Figure 5D-E). Expression of MMP13, which encodes for a key matrix metalloproteinase involved in collagen degradation, denotes active remodeling of the surrounding matrix, while elevated COL10A1 expression reflects the initiation of terminal chondrocyte differentiation and mineralization potential. In contrast, VEGFA expression was comparable between groups (Figure 5F), suggesting that early angiogenic signaling was not substantially influenced by mechanical loading at this stage. When observed under static conditions, dECM-loaded spheroid composites exhibited a trend for increased VEGFA expression compared to control scaffolds (Figure 3E). Together, these findings indicate that short-term dynamic stimulation primarily activates matrix remodeling and hypertrophic differentiation pathways rather than proangiogenic cues.

After 35 d, we evaluated the effects of dynamic compressive loading and RGD concentration on cellular proliferation and early osteogenic activity by quantifying DNA content (Figure 5G) and ALP activity (Figure 5H). Mechanical stimulation resulted in a marked increase in total DNA content, evidenced by a 4.4-fold increase in control scaffolds and a 7.3-fold increase in dECM-loaded spheroid composites compared to their static counterparts (Figure 3B), indicating enhanced cell proliferation under compressive loading. Consistently, ALP activity was significantly elevated in dECM-loaded scaffolds relative to control scaffolds with a 2-fold increase for low RGD (p=0.018) and 1.4-fold increase for high RGD (p =0.04) (Figure 5H), suggesting that the presence of dECM in combination with mechanical cues synergistically promotes endochondral maturation. No significant differences were observed between low and high RGD formulations, indicating that within the tested range, ligand density was less influential compared to the combined effects of mechanical stimulation and dECM bioactivity. The higher expression of Serine Incorporator 5 (Serinc5) (Figure 5I) indicated further commitment to hypertrophic phenotype on dECM constructs compared to controls, regardless of RGD density.(28) This response, when combined with increased vinculin expression, suggests higher cytoskeletal tension that aligns with hypertrophic-osteogenic progression(29). Taken together, these results demonstrate that transient mechanical loading supports cellular expansion within MSC spheroid-laden granular composites and enhances early hypertrophic signaling, reinforcing the mechanobiological advantage of the dECM-loaded alginate granular scaffolds for bone tissue engineering by EO.

Histological analysis of dynamically compressed granular composite scaffolds at d 35 revealed pronounced osteoblastic activity in low RGD-modified constructs containing dECM-loaded spheroids (Figure 6). Enhanced cytoskeletal organization, evidenced by more intense F-actin staining, was observed in dECM-loaded spheroid composites regardless of RGD content. Similarly, we detected more intense OCN staining in dECM-loaded spheroid scaffolds compared to their respective controls. Collagen X staining was undetectable for dECM-loaded scaffolds and high RGD controls, indicating advanced progression of hypertrophic phenotype. Moreover, dECM-laden spheroids in low RGD scaffolds exhibited strikingly higher mineral deposition, exhibited by intense hydroxyapatite staining distributed throughout the scaffold matrix rather than being confined to the spheroid or its peripheral regions (Figure S3).

**Figure 6.**
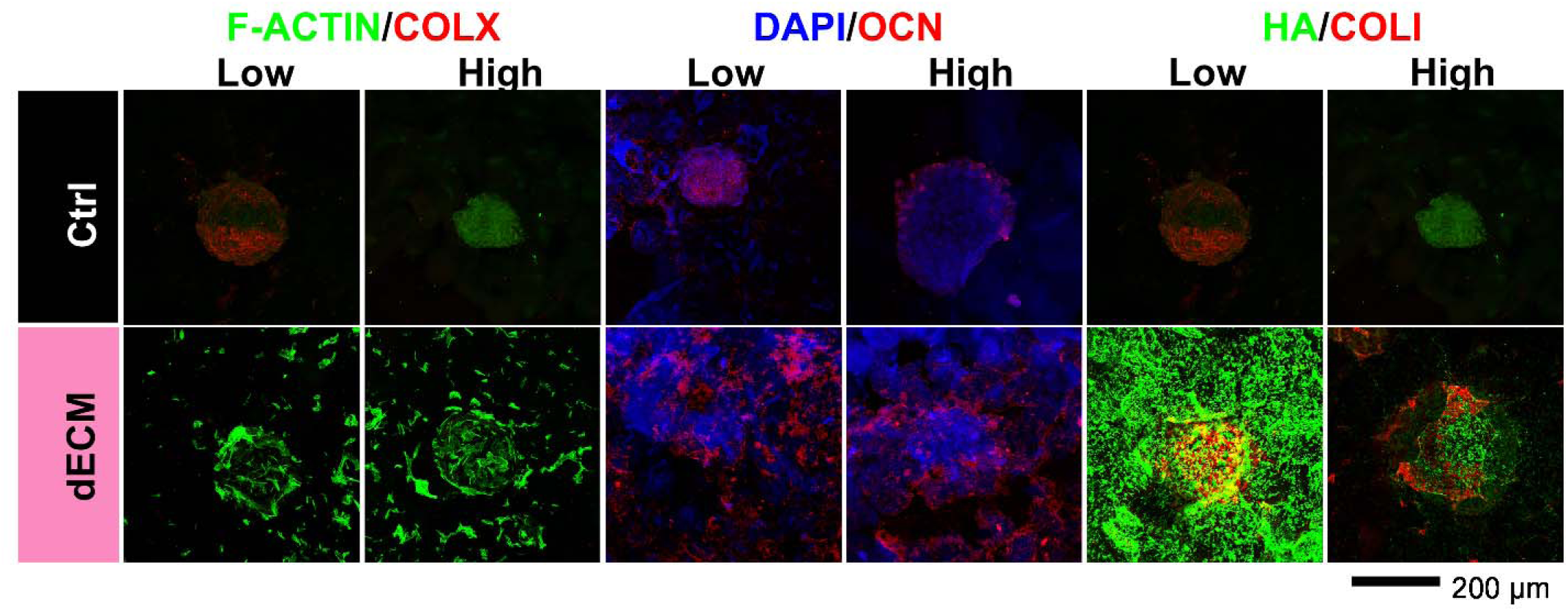
Dynamic compressive loading promotes cell outgrowth, accelerates development of hypertrophic phenotype, and enhances tissue mineralization in dECM-loaded granular composite scaffolds. Histological evaluation of dynamically compressed low and high RGD content dECM-loaded granular scaffolds at d 35. Representative images reveal differences in expression for F-actin, collagen X, cell nuclei, osteocalcin, hydroxyapatite, and collagen I. Scale bar is 200 µm. **Ctrl**, spheroids without dECM; **dECM**, spheroids loaded with dECM; **COLX**, collagen type X; **OCN**, osteocalcin; **HA**, hydroxyapatite**; COLI**, collagen type I.

### Compression-driven hypertrophic maturation is mediated by the Hippo pathway

Our data demonstrate hypertrophic differentiation of dECM-laden MSC spheroids entrapped in low RGD-functionalized alginate microgel scaffolds undergoing dynamic compression. To investigate the mechanism by which compression drives hypertrophic progression for low RGD scaffolds, we used pharmacologic inhibition of the Hippo pathway with verteporfin, a known Yes-associated protein (YAP) antagonist (Figure 7A). The addition of verteporfin during compression results in the downregulation of COLX, MMP13, and VEGFA at d 21 (Figure 7B-D). As expected, verteporfin-treated scaffolds exhibited YAP1 downregulation at d 21 (Figure 7E), whereas untreated scaffolds exhibited upregulation only for dECM-laden scaffolds (Figure 7F). Compared to untreated groups at d 35 (Figure 5G), we observed decreased DNA content of ∼20-fold for control scaffolds and 50-fold for dECM-laden scaffolds (Figure 7G). Although dECM scaffolds treated with verteporfin showed reduction in spheroid sprouting, elevated HA staining for dECM scaffolds suggests a recovery of their mineralization capacity by d 35, indicating a temporal role of YAP/TAZ to coordinate hypertrophic maturation and mineralization (Figure 7H-I). Together, these findings suggest that YAP/TAZ activation is critical to coordinate compressive loading to hypertrophic maturation,

**Figure 7.**
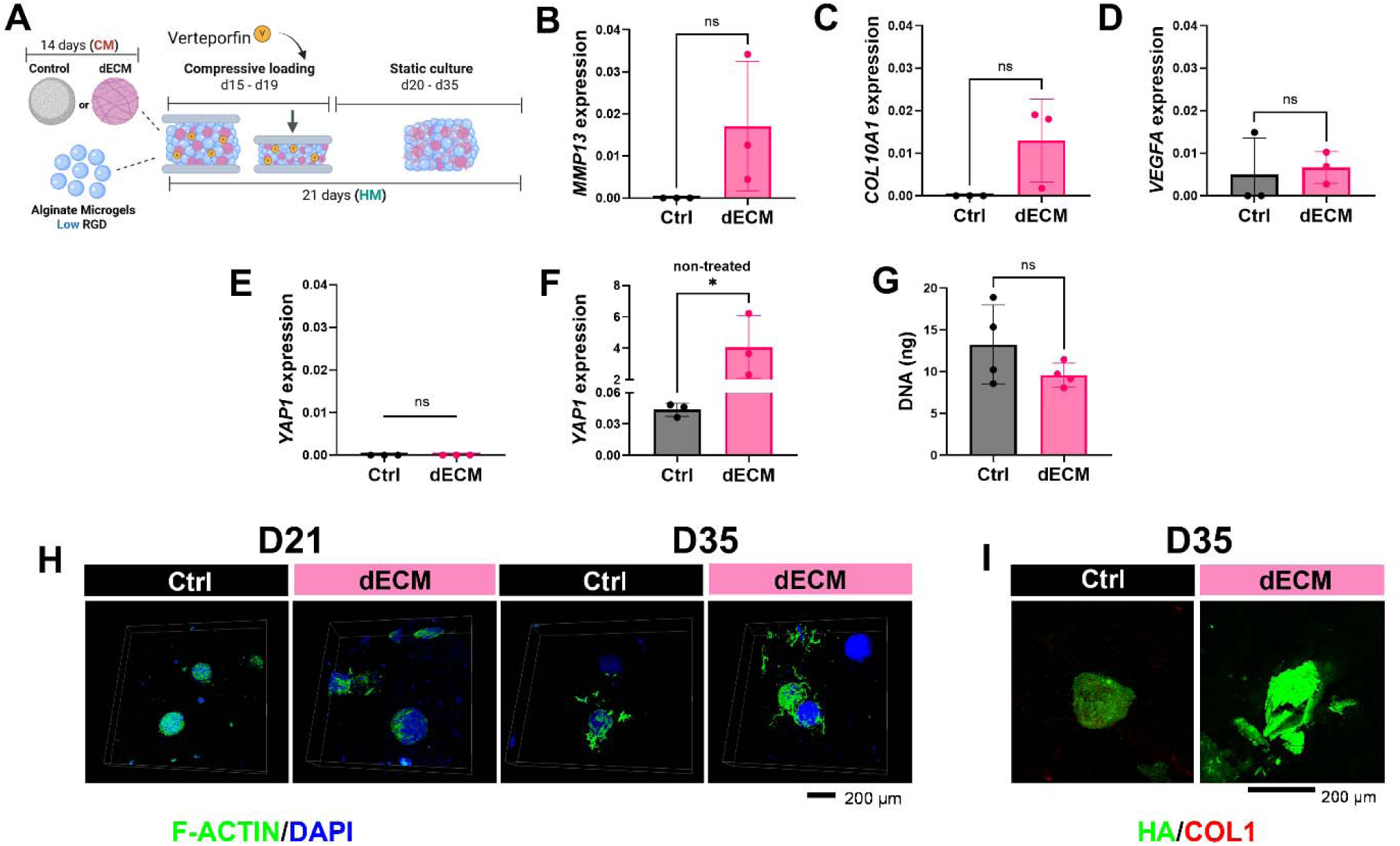
Compression-induced hypertrophic cartilage maturation is associated with YAP/TAZ activation. (**A**) Schematic diagram of fabrication of low RGD alginate granular spheroid-laden scaffolds cultured in chondrogenic conditions for 14 d, followed by dynamic compression for 5 d in the presence of YAP/TAZ inhibitor verteporfin, and cultured in HM under static conditions until d 35. Quantification of gene expression after dynamic compression (d 21) of hypertrophic markers (**B**) *MMP13,* (**C**) *COL10A1,* and (**D**) *VEGFA*. *YAP1* expression for (**E**) treated and (**F**) non-treated scaffolds after compression (d 21). (**G)** DNA content at d 35 indicates changes in proliferation as a result of YAP/TAZ inhibition during compression. (**H**) Fluorescent images of spheroid-laden granular composite scaffolds at d 21 and d 35 treated with verteporfin reveal differences in spheroid sprouting, cytoskeleton architecture (F-actin; green), and cell proliferation (DAPI; blue). (**I**) HA (green) and COL1 (red) staining at d 35 was used to characterize early mineralization. Scale bar is 200 µm. Data are mean ± standard deviation (n=3). *p<0.05. **Ctrl**, spheroids without dECM; **dECM**, spheroids loaded with dECM; **COL1**, collagen type I; **HA**, hydroxyapatite.

We also studied the role of the mechanosensitive ion channel Piezo1 in driving hypertrophic phenotype (Figure S4). Expression of PIEZO1 was upregulated in mechanically stimulated dECM-loaded spheroid composites for both RGD concentrations. The addition of Yoda1, a known Piezo1 agonist, during compression of low RGD constructs resulted in downregulation of MMP13, COLX, and VEGFA at d 21. We also detected a significant decrease in DNA content for control and dECM scaffolds compared to untreated groups (approximately 22-fold) at d 35. Cell staining revealed spheroid sprouting but diminished mineralization by d 35 compared to their static controls or untreated controls. These data suggest that sustained Piezo1 activation alone is insufficient to drive hypertrophic phenotype.

## Discussion

Hypertrophic cartilage engineering leverages maturation of chondrogenic constructs towards a mineralizable, pro-angiogenic phenotype to enable robust endochondral remodeling and bone formation.(3, 30, 31) Periosteum-derived stem cells, bone marrow- or adipose-derived MSCs, and skeletal stem cells are promising sources of chondroprogenitors to generate hypertrophic cartilage,(32, 33), and MSCs are the most studied cell population for cartilage and bone tissue engineering. Under static culture, MSC spheroids embedded within granular microgel scaffolds establish multi-point contacts with surrounding microgels and neighboring spheroids across the inherent interstitial void spaces. This architecture supports short diffusion distances, preserves paracrine signaling, and permits collective behaviors such as cell sprouting, fusion, and ECM deposition that are difficult to achieve in nanoporous bulk hydrogels. Recent reports on granular composite scaffolds similarly demonstrate that spheroid-microgel communication dictates migration and trophic exchange features that map directly onto chondrogenic or hypertrophic tissue assembly.(34) These studies reveal the potential to generate hypertrophic cartilage by applying dynamic compressive loading to composites of MSC spheroids and annealed alginate microgels.

Biomaterials can present instructive biochemical and biophysical cues that promote chondrogenic differentiation by guiding cell adhesion, mechanotransduction, or lineage-specific signaling. Cell-secreted dECM provides a biologically instructive microenvironment that recapitulates many of the biochemical cues present in native cartilage, including collagens, proteoglycans, non-collagenous proteins, and matrix-bound growth factors known to support chondrogenic and hypertrophic progression. By incorporating dECM directly into spheroids, cells are exposed to a rich reservoir of integrin-binding motifs and soluble factors that enhance differentiation through diverse biochemical signals that are absent in traditional synthetic polymers.(35, 36) This is particularly relevant for EO, where the coordinated expression of hypertrophic markers and pro-angiogenic factors such as VEGF is tightly regulated by ECM composition. Moreover, unlike elastic synthetic polymers such as polyethylene glycol or polyvinyl alcohol, alginate’s time-dependent stress relaxation enables cells to dissipate compressive loads, spread, or maintain rounded morphologies.(26, 37) Herein, we used iPSC-derived dECM to address challenges of widespread accessibility and variability. When combined with dECM-loaded spheroids, alginate microgels act as a mechanically tunable yet biologically relevant scaffold, creating an environment where cells can both interpret compressive cues and remodel their microenvironment.

Mechanical loading is a central regulator of chondrogenic differentiation and EO. Compressive cues promote chondrocyte hypertrophy, matrix remodeling, and the transition to a mineralizing template. Compression accelerates hypertrophic differentiation and tissue maturation both in vitro and in vivo, aligning with fracture healing.(3, 38, 39) The macroscale mechanical response of granular hydrogel scaffolds enables enhanced integrin clustering and focal adhesion turnover at cell-microgel interfaces(26), thereby activating loading-responsive signaling cascades that drive hypertrophic differentiation and matrix mineralization. Mechanistically, compressive loading activates integrin/FAK signaling, reorganizes the actin cytoskeleton, and induces YAP/TAZ and Piezo1-mediated Ca²⁺ influx activity toward transcriptional programs that support maturation and ossification.(40) Despite the established importance of mechanical loading in driving endochondral maturation, the effect of these cues and how they are transmitted within granular hydrogel scaffolds was not previously described, motivating our investigation of mechanically induced hypertrophic development in this system.

The balance between adhesion and cytoskeletal tension is well established for regulating MSC differentiation into the osteogenic or chondrogenic phenotypes.(41) However, the influence of RGD ligand density on the development of a hypertrophic chondrocyte phenotype remains poorly characterized, with limited data linking integrin engagement to the chondrocyte-to-hypertrophy transition. Our results suggest a complementary effect of dECM and ligand presentation that modulated hypertrophic differentiation and subsequent mineralization. While lower RGD concentrations promote chondrogenic phenotype for cell aggregates(42), our data suggest that low RGD content induced higher expression of osteogenic markers by d35 in both static and mechanically stimulated conditions, but only with dECM-loaded spheroids. We hypothesize that low RGD content provides a permissive environment for chondrogenic progression into a hypertrophic phenotype by limiting integrin binding compared to high RGD-presenting alginate microgels. This behavior is consistent with prior reports demonstrating that moderate adhesion permits cells to maintain a rounded morphology that supports SOX9-mediated chondrogenesis and cartilage-like matrix deposition.(28)

Prior studies utilizing scrambled RDG peptides as non-adhesive controls confirm poor integrin engagement and attenuation of the Hippo pathway, including YAP/TAZ, resulting in reduced mechanotransducive signaling, cell proliferation, and cytoskeleton-dependent behaviors.(43–45) In contrast, excessive adhesion cues such as those present in high RGD formulations promote integrin clustering, cytoskeletal spreading, and elevated tension, which drives intracellular signaling toward osteogenic commitment in MSCs. In this work, the limited expression of hypertrophic and osteogenic markers in high RGD scaffolds contrasts with reports that an increase in adhesive cues promotes extensive cytoskeletal spreading and tension generation that shifts the balance of intracellular signaling from chondrogenic maintenance toward osteogenic commitment.(46, 47) Importantly, this is the first report of applying dynamic mechanical loading to granular scaffolds. The enhanced responsiveness of dECM-containing spheroids to mechanical cues even after prolonged culture likely results from both biochemical complexity and architectural remodeling within the constructs. The incorporation of dECM introduces a diverse mix of adhesion motifs and matrix-bound growth factors such as TGF-β, BMP-2, and VEGF, which broaden integrin engagement and sustain downstream signaling through mechanoresponsive pathways.(48) Additionally, we hypothesize that dECM integration alters spheroid compaction and microstructure, producing a more interconnected fibrillar network that enhances mechanical continuity between cells and the surrounding matrix. We speculate that this change in organization allows external compressive forces to propagate throughout the spheroid, promoting coordinated mechanotransduction responses. Given that dECM primarily mediates its effects through integrin binding, deformation of the matrix during mechanical loading can dynamically modulate integrin-ligand interactions, thereby altering integrin-dependent signaling pathways and reinforcing load-responsive cellular behavior.(12) Collectively, these biochemical and structural features create a microenvironment that remains mechanically competent even after chondrogenic priming, explaining the sustained mechanosensitivity and hypertrophic progression observed in dECM-loaded granular scaffolds.

At the transcriptional level, dynamic compression upregulated genes associated with matrix remodeling and hypertrophy, such as MMP13 and COL10A1, and activated known mechano-responsive pathways. These findings are consistent with previous reports that cyclic compression can promote chondrogenic or early hypertrophic differentiation or preserve a cartilage-like phenotype depending on the specific loading regime.(39, 49, 50) Consistent with the canonical temporal sequence of EO, dECM-loaded MSC spheroid granular composites exhibited VEGFA upregulation as hypertrophic markers increased. Hypertrophic chondrocytes are a principal source of VEGF in the growth plate, which orchestrates vascular invasion, recruits osteoprogenitors, and coordinates cartilage resorption to bone formation(51). Thus, observing VEGFA upregulation on dECM-loaded scaffolds is concurrent with matrix remodeling and sets the stage for robust vascularization needed for EO-based repair strategies.

While this platform demonstrates the feasibility of dynamically compressed granular scaffolds to create hypertrophic constructs, single-cell and spatial transcriptomic analysis could reveal how local microenvironmental cues and cell-matrix interactions govern hypertrophic development and chondrocyte mineralization. A limitation of this work is the focus on one set of parameters for dynamic compression. In the case of bulk hydrogels, the relation between timing of chondrogenic priming, length of mechanical stimulation, and compression regimen has been studied as a strategy to preserve chondrogenic phenotype.(50) The time-dependent nature of mechanical compression and its effect on dECM-loaded spheroids are yet to be explored for spheroid-laden granular scaffolds. Recent translational studies demonstrate the potential of hypertrophic cartilaginous templates to be rapidly vascularized and remodeled into bone in orthotopic critical-sized defects, supporting the clinical viability of EO-inspired grafts for bone repair(52).

In this work, we present a modular approach to engineering hypertrophic cartilage templates by integrating spheroid-based microgel scaffolds, dECM, and dynamic mechanical compression. Our findings show that macroporous scaffold architecture and compressive loading collectively regulate spheroid-matrix interactions, mechanotransducive signaling, and hypertrophic maturation, while dECM incorporation enhances matrix remodeling and tissue mineralization potential. Importantly, tuning RGD ligand density enabled fine control over adhesion-dependent mechanosensing pathways, highlighting the importance of balanced integrin engagement in guiding hypertrophic progression. Together, these results expand our understanding of hypertrophic chondrocyte development and establish a novel strategy to generate hypertrophic cartilage templates that could yield bioactive, regenerative templates for bone repair.

## Materials and Methods

### Cell culture

Human bone marrow-derived mesenchymal stromal cells (MSCs) (Lot# 00264, RoosterBio, Frederick, MD, USA) from a single donor (female, 19 yo) were expanded and used at passage 5 in standard culture conditions (37°C, 21% O_2_, 5% CO_2_) in growth media (GM) composed of minimum essential alpha medium (α-MEM; Life Technologies; Carlsbad, CA) w/L-glutamine (Invitrogen; Carlsbad, CA) supplemented with 1% penicillin (10,000 U mL^−1^, Gemini; Sacramento, CA) and streptomycin (10 mg mL^−1^, Mediatech; Manassas, VA) (P/S) and 10% fetal bovine serum (FBS; F0926, Lot# 22M276, Sigma-Aldrich). Human MSCs derived from hematopoietic blood-derived iPSCs (iMSC) (ACS-7010; American Type Culture Collection; Manassas, VA, USA) from a single donor (male, 31 yo) were expanded until passage 8-10 in standard culture conditions (37°C, 21% O_2_, 5% CO_2_) in GM. Media for both cell types was refreshed every 2 – 3 d. Chondrogenic media (CM) was composed of Dulbecco’s Modified Eagle Medium (DMEM) (12800-017, Gibco), 10 ng mL^−1^ latent TGF-β1 (R&D systems, Minneapolis, MN), 1 mM sodium pyruvate, 40 μg mL^−1^ L-proline, 1.5 mg mL^−1^, bovine serum albumin (BSA), 1x insulin-transferrin-selenium (ITS), 50 nM dexamethasone (DEX), and 50 µg mL^−1^ L-ascorbic acid-2-phosphate (A2P) (all from Millipore Sigma, Burlington, MA), and 1% P/S. Chondrogenic conditioning was carried out in hypoxic culture conditions (37°C, 5% O_2_, 5% CO_2_) attained in a Heracell 150i tri-gas incubator (Thermo Electron, Asheville, NC). Hypertrophic media (HM) consisting of DMEM, 1% FBS, 50 µg mL^−1^ A2P, 10 nM DEX, 7 mM of β-glycerophosphate, and 1 nM triiodothyronine was used to induce chondrocyte differentiation after 14 d in chondrogenic conditions culture. Hypertrophic differentiation was conducted in standard culture conditions (37°C, 21% O_2_, 5% CO_2_) for 21 d.

### Decellularization of cell-secreted ECM

Decellularized cell-secreted ECM (dECM) from iMSCs was produced in standard culture conditions (37°C, 21% O_2_, 5% CO_2_) via macromolecular crowding as we described(*16, 19*). Briefly, iMSCs were cultured in GM supplemented with A2P (50 μg mL^−1^) and expanded until 80% confluence. After preconditioning, cells were seeded at 50,000 cells cm^−2^ in the presence of a macromolecular crowding agent (λ-carrageenan, 75 μg mL^−1^, 22049; MilliporeSigma) supplemented with A2P for 9 d. ECM was decellularized using an enzymatic method consisting of 0.1 mg mL^−1^ DNAse I (MilliporeSigma) for 1 h at 37°C to yield dECM. After PBS rinsing, dECM was scraped, collected in sterile 0.02 N acetic acid, sonicated to produce a homogeneous solution, quantified, and stored at −20°C until use.

### Spheroid formation and differentiation

Suspension plates (24-well) containing 29 agarose microwells per well were used to make spheroids of 5,000 MSCs with or without dECM. After trypsinization, 1 mL of cell suspension was pipetted into each well at a concentration of 1.45×10^5^ cells mL^−1^, followed by centrifugation at 900 x g for 8 min. For dECM-loaded spheroids, dECM was mixed within the cell suspension at a concentration of 48.4 µg mL^−1^ to achieve a concentration of 1.67 µg per spheroid (following our work of 0.333 ng of dECM per cell) and centrifuged as described(*19, 20*). Spheroids were cultured in GM in static conditions for 48 h to allow cell aggregation. After aggregation, spheroids were cultured in CM under hypoxic conditions (37°C, 5% O_2_, 5% CO_2_) for 14 d to induce chondrogenic differentiation. After chondrogenic conditioning, spheroids were cultured in HM for 21 d in standard culture conditions (37°C, 21% O_2_, 5% CO_2_). Media was refreshed every 2-3 d. Spheroid diameter was quantified using brightfield micrographs and ImageJ (NIH, Bethesda, MD). DNA content was quantified by a Quant-iT^TM^ PicoGreen assay. Glycosaminoglycan (GAG) content in the spheroids was measured *via* the Dimethylmethylene Blue (DMMB) assay. ALP activity was quantified *via* hydrolysis of p-Nitrophenyl phosphate (pNPP) substrate to study changes in osteoblastic activity and tissue mineralization as part of maturation of hypertrophic phenotype.

### Alginate functionalization

Prior to microgel fabrication, alginate was modified with RGD peptide and 2-aminoethylmethacrylate (AEMA) to facilitate cell adhesion and UV-crosslinking, respectively. Alginate (PRONOVA UP VLVG alginate; < 75,000 g mol^−1^; NovaMatrix, Sandvika, Norway) underwent methacrylation (22.5% theoretical methacrylation) as described(53). Briefly, 2 g of alginate was dissolved in 1% (w/v) 50 mM (2-(N-morpholino)ethanesulfonic acid (MES) and mixed with 0.265 g of N-hydroxysulfosuccinimide sodium salt (sulfo-NHS) and 0.875 g of 1-ethyl-3-(3-dimethylaminopropyl)-carbodiimide hydrochloride (EDC) to activate the carboxylic acid groups of the alginate. After 5 min, 0.360 g of AEMA was added (molar ratio of NHS:EDC:AEMA = 1:2:1), and the reaction was maintained at room temperature for 24 h. Afterwards, the reaction mixture was precipitated in cold acetone and redissolved in ultrapure water. The resulting solution was dialyzed against distilled water for 72 h using 3500 Da MWCO dialysis membrane (Spectrum Laboratories, New Brunswick, NJ), with water changes every 6-24 h. The dialyzed alginate was lyophilized. Alginate was further functionalized with Arg-Gly-Asp (RGD) peptide through carbodiimide-mediated coupling reaction. Briefly, methacrylated alginate (MEA) was dissolved at 1% (w/v) in 50 mM MES buffer and allowed to equilibrate overnight at RT. The next day, EDC and sulfo-NHS were added at a 2:1 ratio per gram of alginate. The peptide G₄RGDSP (GenScript, Piscataway, NJ) was added in various amounts to achieve a degree of substitution (DS) of 2 (low) or 20 (high) per alginate polymer chain. The reaction product was dialyzed (3.5 kDa MWCO; Spectrum Laboratories) against ultrapure water for 3 d with regular water changes, sterile filtered, and lyophilized. Alginate was protected from light and stored at −20°C until used.

### Fabrication of alginate microgels

To fabricate microgels, we used a multichannel microfluidic device design first reported by de Rutte et al.(7) and since adapted by our group.(8, 9) Low and high RGD-functionalized VLVG alginate dissolved in 100 mM Ca-EDTA at 2% (w/v) comprised the dispersed phase, while 0.5% (v/v) Picosurf (SphereBio, Cambridge, UK) in NovecTM 7500 (3MTM, Maplewood, MN) comprised the continuous phase. Microgels were collected off device in a 1% (v/v) acetic acid solution. Upon reduction in pH, calcium ions disassociate from EDTA, enabling ionic crosslinking of alginate chains. An equal volume of 20% (v/v) Perfluoro-1-octonal (MilliporeSigma) in NovecTM 7500 was added to crosslinked microgels to destabilize the emulsion. Microgels were subsequently rinsed with 70% isopropyl alcohol (x3) and sterile distilled water (x3) to remove remaining oil. To form annealed scaffolds, microgels were jammed at 14,000xg in a 0.3% (w/v) IrgaCure (MilliporeSigma) solution in cell culture media. Scaffolds were exposed to UV light (320-500nm, OmniCure S2000, Excelitas Technologies, Mississauga ON, Canada) at 20 mW cm^−2^ for 5 min.

### Characterization of alginate microgels

Microgels diluted in DI water were pipetted onto a glass-bottom petri dish and imaged at 10x using an EVOS microscope. Microgel diameter was manually measured using ImageJ with at least 3 fields of view. Compressive moduli of individual microgels were measured using a MicroTester (CellScale, Waterloo, ON)(8). Microgels were compressed over 30 s to 30% of their original diameter by a 0.5 x. 0.5 mm stainless-steel platen attached to a 0.1524 mm diameter tungsten rod. The displacement and force of each microgel were recorded and utilized to calculate the modulus. The modulus was calculated from the linear region of the compressive modulus versus the nominal strain graph using a custom Python code.

Annealed microgel scaffolds were placed in 300 µL of 1 mg mL^−1^ of FITC-Dextran (200 kDa, Sigma) overnight at RT. Samples were washed three times with DI water and placed on a glass-bottom petri dish to image on a confocal microscope. The z-stacks had a step size of 2.5 µm and a total height of 300 μm. Pore volume and total volume were calculated to determine the porosity using IMARIS (Oxford Instruments)(23). We measured the compressive moduli of annealed microgel scaffolds using an Instron 3345 Compressive Testing System (Norwood, MA). Hydrogels were measured following 24h in cell culture medium. Hydrogels were loaded between two flat platens and compressed at a rate of 0.05 mm s^−1^. Moduli were calculated from the slope of stress versus strain plots limited to the linear first 10% of strain.

### Fabrication of dECM-loaded spheroid-laden alginate microgel composite scaffolds

Spheroids with and without dECM were cultured in chondrogenic conditions as previously described.(19) After 14 d of chondrogenic priming, spheroids were collected and mixed with alginate microgels to form alginate microgel dECM-loaded spheroids granular scaffolds. Based on the original spheroid cell density, 44 spheroids were mixed with microgels to fabricate a 4 mm cylindrical scaffold with a final volume of 22 μL and a cell density of 10×10^6^ cells mL^−1^. The mixture of microgels and spheroids was annealed into a composite as previously described by exposure to UV light (20 mW cm^−2^) for 5 min to anneal and transferred to a non-adhering tissue culture plate. Scaffolds were placed in HM, and media was changed every 2-3 d and cultured under standard cell culture conditions.

### Compressive loading of spheroid-laden alginate microgel composite hydrogels

Granular scaffolds of 5 mm diameter were fabricated as described above and allowed to equilibrate in HM media for 24 h before mechanical stimulation. Briefly, 58 spheroids were collected after 14 d of chondrogenic priming and mixed with 30 µL of either low or high RGD alginate microgels to form composite scaffolds with a cell density of 10×106 cells mL^−1^. Immediately after fabrication, scaffolds were placed in a non-adherent tissue culture plate and left in HM to equilibrate for 24 h. The following day, scaffolds were transferred into BioFlex® 6-well plates and moved to a Flexcell® compression bioreactor (FX-5000™ Compression System). The scaffolds were compressed to a strain of 25% at a frequency of 1 Hz for a period of 2 h, followed by a rest time of 10 h. The compression regimen was applied for 5 consecutive days, and scaffolds were transferred to a non-adherent tissue culture plate for the remainder of the experiment. Scaffolds were maintained in HM in standard cell culture conditions and collected after 7 and 21 d from the start of the compressive loading regimen.

To study the mechanistic effects of compression on hypertrophic differentiation, we supplemented HM with either YAP/TAZ inhibitor verteporfin (5 µM; Sigma, SML0534), or Piezo1 agonist Yoda1 (5 µM; Sigma, SML1558) during the 5 d compression regime. After a 48 h resting period, scaffolds were either collected (d21) or maintained in static conditions in HM for 2 additional weeks (d35).

### Gene expression

Total RNA was isolated from cells using TRIzol reagent (ThermoFisher) according to the manufacturer’s instructions. RNA quality and quantity were measured using a Nanodrop One instrument (ThermoFisher) before reverse transcribing to cDNA with the QuantiTect Reverse Transcription kit (Qiagen, Hilden, Germany). All cDNA samples were diluted with PCR-grade ultrapure water to 12.5 ng μL-1 prior to qPCR. qPCR was performed using a Taq PCR Master Mix kit (Qiagen), TaqMan Gene Expression Assay probes (cat. 4331182, ThermoFisher), and a QuantStudio 6 instrument (ThermoFisher). Samples were activated at 94 °C for 3 min, followed by 40 cycles of 94 °C for 30 s, 60 °C for 30 s, and 72 °C for 1 min, and underwent a final annealing step at 72 °C for 10 min.

Expression of chondrogenic differentiation markers transcription factor SOX-9 (SOX9, Hs00165814_m1) and aggrecan (ACAN, Hs00153936_m1) were interrogated at d 14. The expression of hypertrophic phenotype markers matrix metalloproteinase 13 (MMP13, Hs00942584_m1), collagen type X (COL10A1, Hs00166657_m1), and vascular endothelial growth factor A (VEGFA, Hs00900055_m1) were evaluated at d 21 and 35. Gene expression for osteogenic markers collagen 1 (COL1A1, Hs00164004_m1) and osteopontin (SPP1, Hs00959010_m1) was assessed at d 35. Expression of mechanosensitive markers Piezo1 (PIEZO1, Hs00207230_m1) and Yes-associated protein 1 (YAP1, Hs00902712_g1) were used to interrogate the effects of dynamic compression. Gene expression was normalized housekeeping gene GAPDH (Hs02786624_g1) to yield ΔCt. Freshly formed spheroids without dECM were used as a control to calculate ΔΔCt. Fold change was calculated using the 2−ΔΔCt method.

### Histological evaluation

Single spheroids and spheroid-laden granular scaffolds were fixed in 4% paraformaldehyde for 1 h at room temperature, rinsed with PBS, and stored at 4°C for further histological evaluation. For single spheroids, samples were collected and embedded in Histogel (ThermoFisher), dehydrated, paraffin-embedded, and sectioned at 7-8 µm thickness. Sulfated glycosaminoglycans (GAGs) and calcium deposits were visualized using Safranin-O and Alizarin Red staining, respectively. For immunohistochemistry, paraffin was removed by heating at 60°C for 10 min, followed by gradual rehydration and antigen retrieval in citrate buffer. After PBS washes, sections were blocked for 45 min at room temperature with 10% goat serum in 1% BSA and incubated overnight at 4°C with primary antibodies against collagen II (1:200, Abcam ab34712), SOX9 (1:200, Abcam ab76997), collagen X (1:200, Abcam ab58632), serinc5 (1:200, Abcam ab204400), or vinculin (1:200, ThermoFisher 14-9777-82) diluted in 0.1% BSA. Following PBS washes, slides were incubated for 1 h at room temperature with either Alexa Fluor 568–conjugated goat anti-mouse IgG (1:600, Abcam ab175473) or Alexa Fluor 647–conjugated goat anti-rabbit IgG (1:600, Abcam ab150083) in 0.1% BSA.

Spheroid-laden granular scaffolds were embedded in 4% low-melting-point agarose (A9414, Sigma) to preserve scaffold structure. After cryoembedding in optimal cutting temperature (OCT) media, samples were sectioned at 11-12 µm thickness. Sections were blocked and stained for collagen X (1:200, ThermoFisher PA5-115039), collagen I (1:200, Abcam ab138492), and osteocalcin (1:200, Abcam ab93876), similar to paraffin-embedded samples. Bulk scaffolds were stained with DAPI (4’,6-diamidino-2-phenylindole, ThermoFisher) and phalloidin (A12379, ThermoFisher) to visualize spheroid sprouting and cellular ingrowth. OsteoimageTM (PA-1503, Lonza) was used to assess hydroxyapatite secretion within histological sections and bulk scaffolds. Imaging was performed at 20× and 40× magnification using a Leica Stellaris 5 confocal microscope.

### Statistical analysis

All experiments were repeated independently at least three times and presented as mean ± standard deviation. Statistical analysis was performed using Prism 10.2.3 software (GraphPad, San Diego, CA) utilizing one-way or two-way ANOVA or an unpaired Student’s t-test as applicable for multiple or pairwise comparisons, respectively. Comparisons between each condition were evaluated using Pearson’s correlation coefficient (two-way ANOVA) or post-hoc Tukey test (one-way ANOVA). Groups with different letters indicate statistically significant differences (p<0.05), while groups with the same letters are not significant.

## Acknowledgments

We thank Mason Caserta, Connor J. Dorais, and Advait Bhagvat for their technical assistance that greatly contributed to the completion of this work.

## Author contributions

Conceptualization: JKL, DHRR

Methodology: DHRR, ACF, SRP, SWF, EEW

Investigation: DHRR, ACF, SWF

Visualization: DHRR, JKL

Supervision: JKL

Writing—original draft: DHRR, JKL

Writing—review & editing: DHRR, ACF, SRP, SWF, EEW, JKL

## Data and materials availability

All data are available in the main text or the supplementary materials. Additional data related to this paper may be requested from the authors.

## Supplementary data

**Fig. S1.**
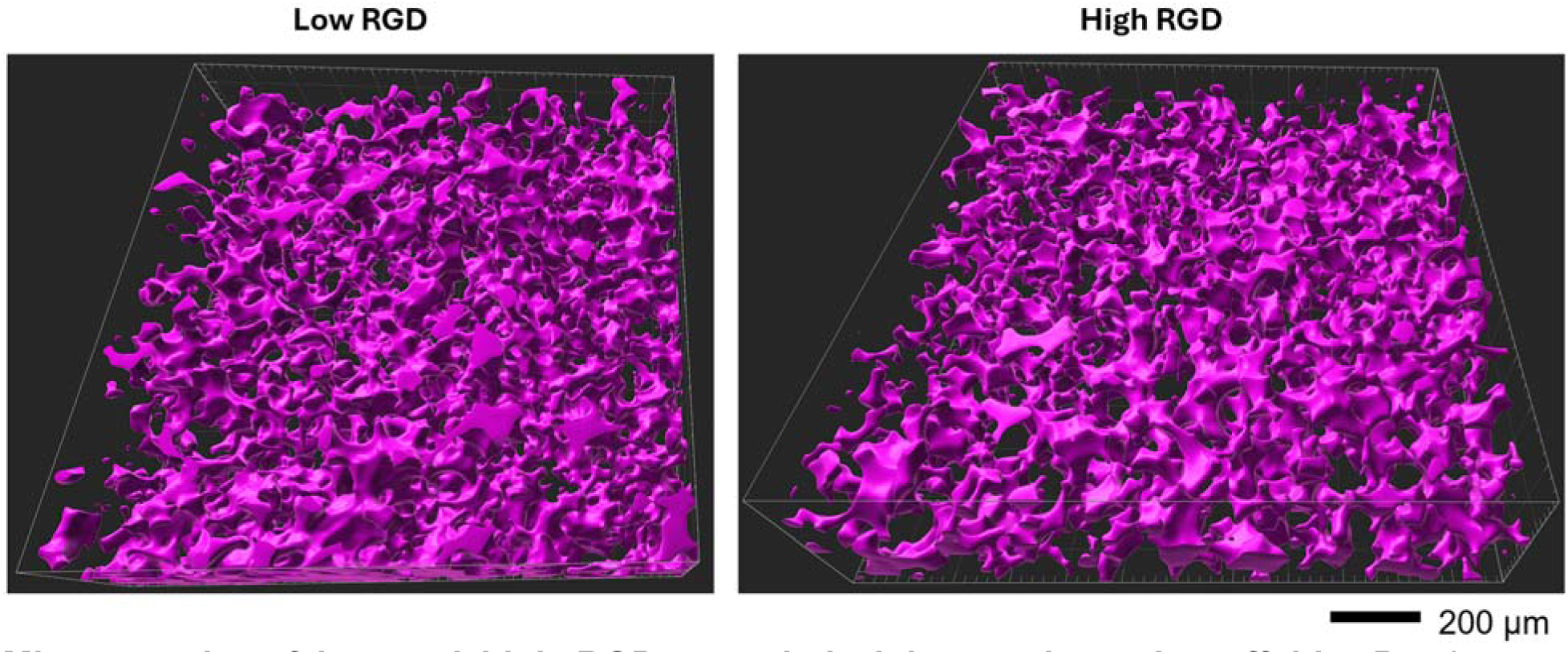
Microporosity of low and high RGD annealed alginate microgel scaffolds. Purple area denotes void space occupied by FITC-dextran. Fluorescent 3D micrographs were rendered and analyzed using IMARIS. Scale bar is 200 µm.

**Fig. S2.**
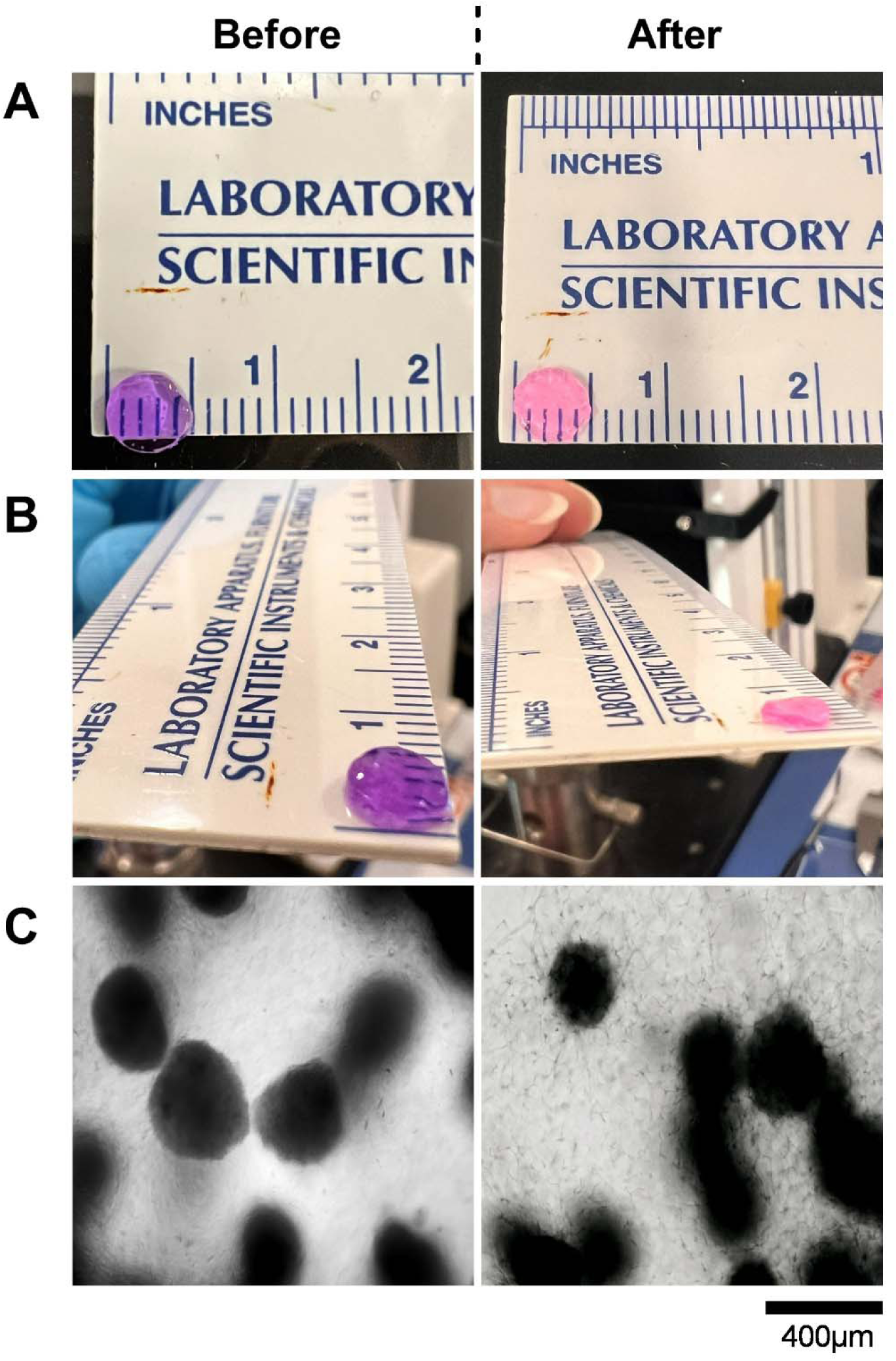
Representative images of spheroid-laden microgel scaffolds (low RGD) before and after dynamic compression. Gross images show no changes in **(A)** scaffold diameter (5 mm) but minimal change in **(B)** thickness (∼1mm). **(C)** Brightfield images highlight the effects of dynamic compression on spheroid sprouting. Scale bar is 400 mm.

**Fig. S3.**
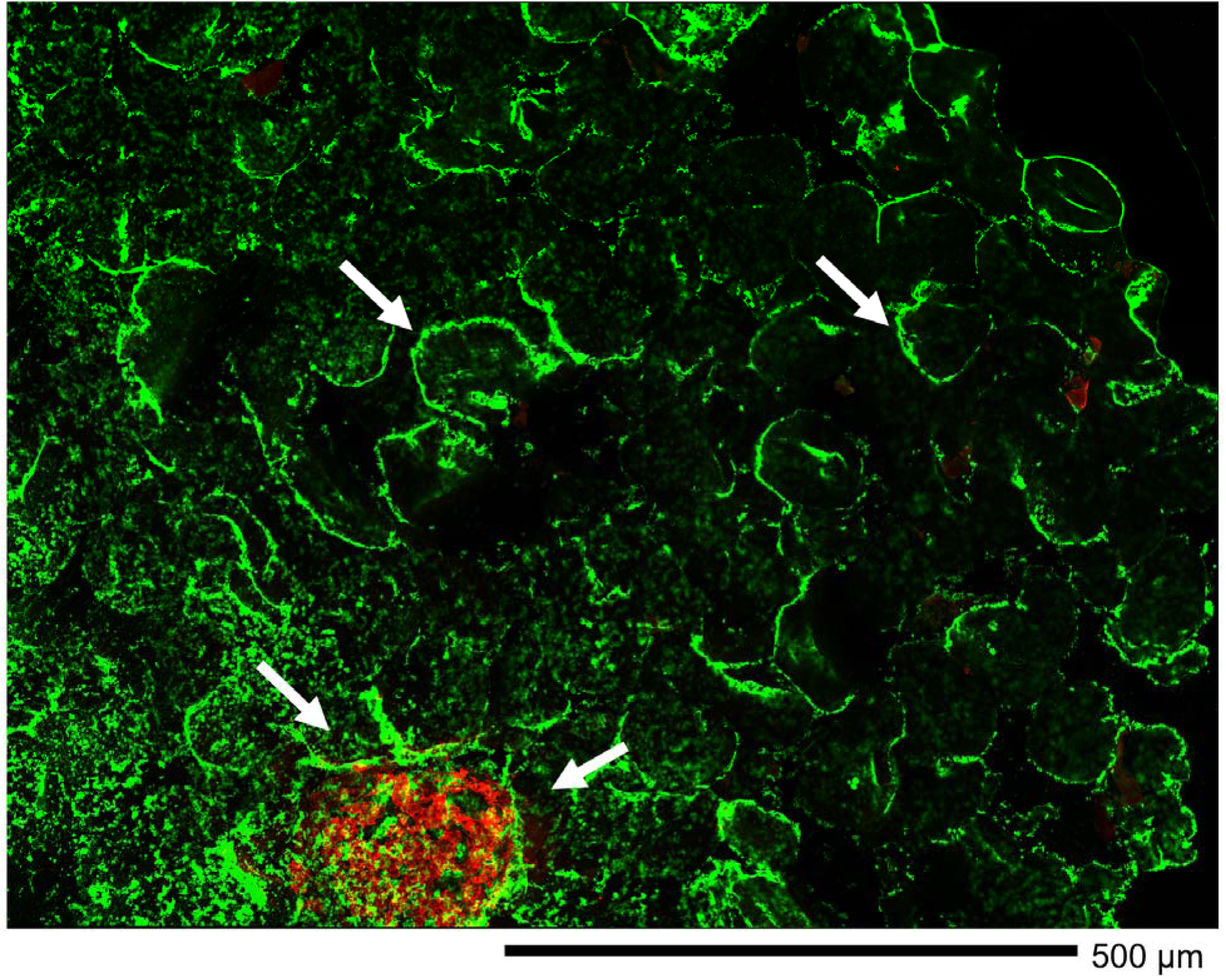
Representative image of a mechanically stimulated alginate microgel (low RGD) dECM-laden spheroid scaffold at d 35. Hydroxyapatite (green) and F-actin (red) staining show extensive mineralization across a large section of the scaffold. Mineralization is observed both adjacent to the spheroids and along the periphery of the microgels (white arrows). Scale bar is 500 µm.

**Fig. S4.**
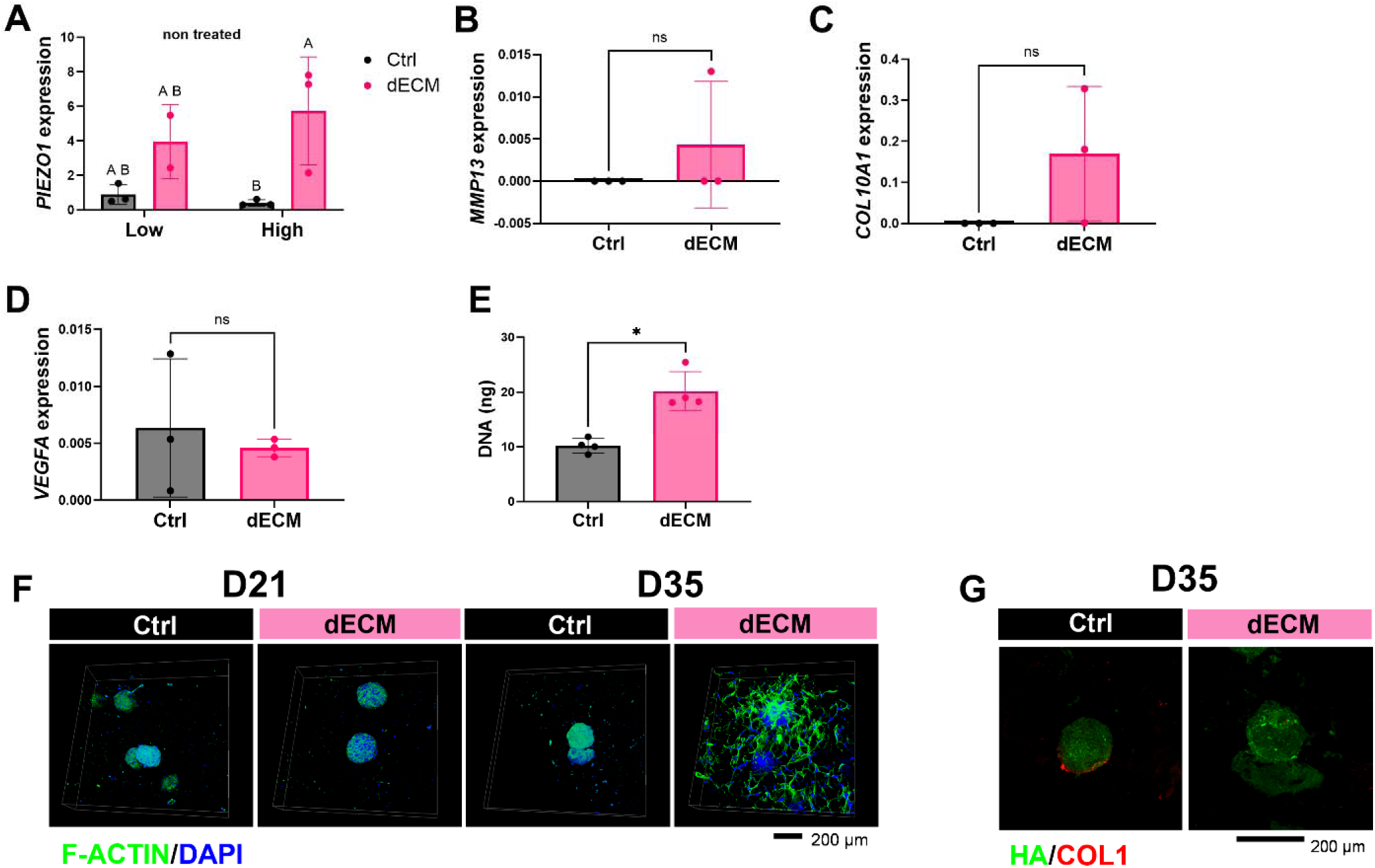
Chemical activation of Piezo1 is not associated with maturation of compression-induced hypertrophic cartilage. (**A**) Quantification of *PIEZO1* in non-treated dynamically compressed scaffolds shows upregulation for dECM-laden spheroid scaffolds regardless of RGD content. Activation of Piezo1 during compression using Yoda1 reveals downregulation of hypertrophic markers (**B**) *MMP13*, (**C**) *COL10A1*, and (**D**) *VEGFA.* (**E)** DNA content at d 35 indicates changes in proliferation because of Piezo1 continuous activation during compression. (**F**) Fluorescent images of spheroid-laden granular composite scaffolds at d 21 and d 35 treated with Yoda1 reveal differences in spheroid sprouting, cytoskeleton architecture (F-actin; green), and cell proliferation (DAPI; blue). (**G**) HA (green) and COL1 (red) staining at d 35 was used to characterize early mineralization. Scale bar is 200 µm. Data are mean ± standard deviation (n=3). *p<0.05. **Ctrl**, spheroids without dECM; **dECM**, spheroids loaded with dECM; **COL1**, collagen type I; **HA**, hydroxyapatite.

